# Finding space for rewilding: participatory scenarios reveal ecological opportunities based on plural values of nature

**DOI:** 10.1101/2025.03.05.641619

**Authors:** Laura C. Quintero-Uribe, Henrique M. Pereira, Rowan Dunn-Capper, Jenny Schmidt, Néstor Fernández

## Abstract

1. Rewilding aims to restore degraded ecosystems in human-dominated areas by promoting self-sustained ecosystems and reducing human control over ecological processes. However, co-benefits and trade-offs between rewilding actions and the interests of local societies must be thoroughly considered. IPBES proposed the Nature Futures framework, a scenario tool to engage society in conservation planning. However, its potential for co-designing ecological restoration remains to be explored.
2. This paper presents a novel methodology that utilises the Nature Futures Framework to address opportunities, co-benefits and trade-offs using rewilding. The study aims to investigate the Nature futures Framework effectiveness in rewilding by involving stakeholders in participatory scenario development. To accomplish this, we conducted stakeholder interviews to co-design three distinct scenarios that reflect diverse values of nature and rewilding strategies for the German Oder Delta.
3. The scenario development process identified spatial priorities for three scenarios: Nature for Nature, Nature for Society (NfS) and Nature as Culture. (NaC). The first, focused on rewilding and increasing forest and river connectivity across 65.8% of the study area. NfS narratives covering 14.4% of the area emphasised de-intensification of forestry and reflooding of peatlands, while NaC prioritised transformation towards extensive agriculture and reflooding for paludiculture across 8.6%.
4. The study identified regions with single and multiple values of nature. Further, there were regions of overlap between scenarios, highlighting the interconnectedness of different rewilding actions and priorities. By pinpointing corresponding values, the research highlighted actions that could result in co-benefits. The findings emphasise the importance of considering pluralistic values when designing rewilding actions, which encourages stakeholders to highlight areas of less resistance to rewilding and co-adapt to the changes in the landscape.
5. This study presents a methodological framework for participatory scenario development for rewilding initiatives. By combining participatory processes with the Nature Futures Framework, our structured approach helps identify priority areas and navigate socio-ecological challenges in landscape restoration. It employs participatory mapping to aid the visualization of land-use changes and aligning rewilding actions with ecological and societal needs. This approach enhances governance frameworks and promotes inclusive rewilding efforts, creating a replicable model for landscape restoration that connects rewilding with sustainability and societal well-being goals.

## 1 INTRODUCTION

Rewilding has emerged as a pragmatic restoration strategy aimed at reverting biodiversity loss and ecosystem degradation while seeking co-benefits for local societies (Fernández et al., 2017; Perino et al., 2022; Svenning, 2020). It addresses the pressing need to address human-induced ecosystem degradation and its links to human well-being, highlighting the urgent requirement for transformative change (Massenberg et al., 2022). The European Green Deal (European Commission, 2019) and the Kunming-Montreal Global Biodiversity Framework (Convention on Biological Diversity, 2022) underscore the importance of sustainable development and ecosystem restoration. However, a notable gap persists in translating these policies into practical actions due to a lack of tools to better navigate the complex interrelation between nature and society (IPBES 2016).

To navigate this gap, envisioning future scenarios becomes a transformative approach, reframing the relationship between people and nature. However, existing envisioning tools like the Shared Socio-economic Pathways and climate change scenarios (SSPS) have limitations in recognizing the diverse interconnections and feedback loops between nature and people (IPBES, 2016). In this context, rewilding, as an approach embedded in ecological restoration science, offers a unique opportunity to address these limitations (Fernández et al., 2017; Perino et al., 2019).

The Nature Futures Framework (NFF) stands out as a crucial tool that aligns with rewilding goals and can enhance its implementation. By incorporating diverse value perspectives and employing participatory methods such as Participatory Scenario Development, the NFF offers a novel and participatory approach to exploring the complexities of socio-ecological systems (Lundquist et al., 2021; Pereira et al., 2020). The framework guides spatial planning and facilitates the exploration of policy consequences, identifying strategies for sustainable futures in biodiversity and ecosystems (Kim et al., 2023).

By integrating the NFF with the pressing need to create and implement actions to halt biodiversity loss, a promising synergy addresses the challenges of rewilding at both theoretical and practical levels. This integrated approach ensures that rewilding initiatives consider the complex interconnected socio-ecological factors at various scales, fostering a harmonious relationship between nature and society (Dunn-Capper et al., 2023; Perino et al., 2019; Van Meerbeek et al., 2019). The NFF, with its participatory and co-design principles, becomes a valuable tool in envisioning and implementing rewilding projects that maximize co-benefits, navigate trade-offs, and contribute to transformative change towards sustainable development while halting biodiversity loss (Durán et al., 2023). Prior attempts to co-design restoration and conservation initiatives often failed to consider the varied value perspectives crucial for successful biodiversity restoration (Quintero-Uribe et al., 2022). To our knowledge, this study is the first to utilize the NFF in developing scenarios for rewilding.

Our primary research goal is to explore how developing participatory scenarios that envision diverse nature futures can help uncover co-benefits and trade-offs in rewilding practices. We have chosen the German Oder Delta as our study system because of its rich socio-ecological context and unique conservation challenges. By actively engaging with stakeholders and applying the Nature Futures Framework (NFF), we aim to assess how pluralistic value perspectives impact rewilding activities in this complex landscape. Additionally, our study aims to identify co-benefits and trade-offs associated with rewilding practices by synthesizing restoration narratives and incorporating local knowledge through participatory scenario development. By emphasizing the significance of the Oder Delta’s intricate socio-ecological dynamics, our research seeks to provide nuanced insights into the values of nature and the potential impacts or benefits of rewilding actions. This approach contributes to the advancement of rewilding practices and enhances our understanding of effective landscape restoration strategies in the European context.

## 2 METHODS

### 2.1 Study area

The Oder Delta region is situated on the Polish-German border, located at the mouth of the Oder River as it empties into the Baltic Sea (Fig 1). The region includes a variety of habitats, including peatlands, marshes, swamps, wet meadows, and forests, and is home to numerous species of birds, fish, and mammals. The Ueckermunde Heide region features sandy and dune landscapes, with diverse land use including arable farming, grassland farming, and forestry. Despite the challenge of poor soils in floodplain areas, natural resources play a crucial role in rural development, providing goods and services such as wood and foods (Beetz et al., 2007; Hirschfeld et al., 2009). Efforts to transition to mixed forests and the presence of large competitive farms alongside smaller organic farms highlight the region’s evolving agricultural landscape. In the region, wheat cultivation on fertile soils and livestock farming, particularly cattle and pig farming, are significant, with a fifth of the agricultural land dedicated to permanent grassland (Beetz et al., 2007; Plieninger et al., 2007).

**FIGURE 1.**
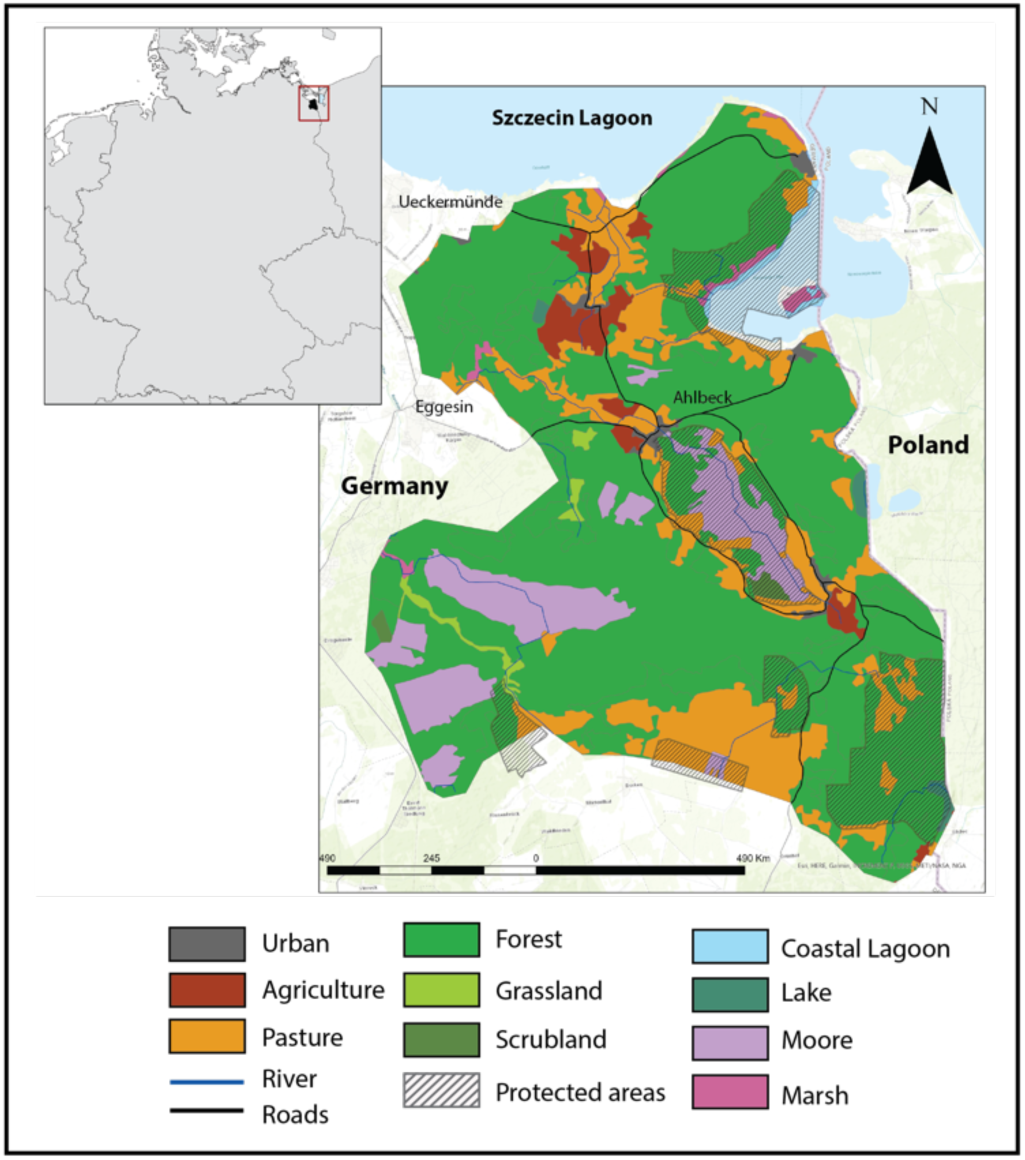
Map of Uckermünde Heath region in the Oder Delta region and the main land uses (CORINE Land Cover, 2018)

Today, the region is a popular ecotourism spot, offering activities such as birdwatching, hiking, and swimming. Furthermore, it is an ecologically important site, providing crucial habitat and breeding grounds for various species and serving as a critical biodiversity hotspot for the region’s inland ecosystems (German Federal Environmental Foundation, 2023).

### 2.2 Participatory scenario development process

Rewilding practitioners were involved in the research process to ensure robust and relevant results. Stakeholder groups were included in scoping and planning the Oder Delta rewilding. As a starting point, we interviewed officials from the Rewilding Oder Delta e.V. (ROD) organization and expert researchers to understand the region’s situation, challenges, opportunities, and regulations. We carried out the scenario development process in three phases: 1) Stakeholder’s analysis for the identification and selection of the contributing stakeholders; 2) Development of the overall Nature Futures scenario storylines through semi-quantitative interviews and participatory mapping of the scenarios; and 3) Data analysis through Systematic content analysis of the interviews and the digitalization of participatory maps (Fig S1).

### 2.3 Participant identification and sampling

We conducted stakeholder identification and interviews from September 2021 to March 2022 through online video calls, required by COVID-19 travel restrictions. To address challenges such as limited stakeholder availability and a tight project timeline, we selected a small group of knowledgeable individuals in biodiversity conservation and sustainable land management. Using participatory mapping methods and the Net-Map tool, we identified key actors and evaluated their connections and influence (Schiffer & Hauck, 2010). Collaborating with officials from Rewilding Oder Delta e.V. (ROD), we aimed to identify a diverse group of stakeholders actively engaged in local decision-making, thereby providing rich insights into the complex socio-ecological dynamics of the community.

From our analysis, we selected nine key stakeholders from agriculture, forestry, nature conservation, and administrative institutions for interviews, concluding once we reached the principle of saturation point where additional interviews no longer yielded new information. While a limited participant pool can restrict diverse perspectives potentially resulting in biased outcomes (Pahl-Wostl, 2002), it has been demonstrated that small group of stakeholders can yield valuable qualitative insights, especially in early project stages (Dunn-Capper et al., 2023; Rohrbach et al., 2016).

By strategically selecting participants and employing rigorous and structured methodologies (PPGIS), we ensured the reliability of the information gathered. This focus on depth over breadth allowed for a nuanced exploration of potential futures in rewilding, capturing the region’s complex ecological and social dynamics (Quimby & Beresford, 2023; Rohrbach et al., 2016). Our findings demonstrate that carefully designed participatory mapping and respondent-driven sampling can produce reliable data.

Ethics approval was granted by the lead supervisor in accordance with university guidelines. The study adhered to the German Research Centre for Integrative Biodiversity Research’s code of conduct and GDPR regulations. Informed consent was obtained from all stakeholders, ensuring confidentiality and data protection. A consultancy firm supported participatory processes ensuring that all necessary steps were taken to avoid putting stakeholders at risk or in vulnerable positions. To protect participants discussing sensitive topics like rewilding, interviews were fully anonymized, and raw data was securely stored in password-protected folders accessible only to principal investigators.

### 2.4 Development of Scenario storylines

We developed the initial scenario storylines by employing the methodology established in Quintero-Uribe et al. (2022), which maps the various elements of Rewilding and the contributions of Nature to human well-being. These storylines will be presented to the final selection of stakeholders chosen for the interviews. It is important to note that the scenario storylines were constructed prior to conducting the interviews.

We utilised the NFF and its different nature value perspectives as a guide to develop three overall scenario storylines for rewilding landscapes in the Oder Delta. The Nature Futures Framework (NFF) is a tool for envisioning positive outcomes for nature and people. The NFF delineates three primary axes of values for nature and people: Nature for Nature (intrinsic values), Nature for Society (instrumental values), and Nature as Culture (relational values) that represent people’s preferences for the future of nature (Kim et al., 2023; Pereira et al., 2020).

This study integrates the three NFF value perspectives with Perino et al.’s (2019) rewilding framework—trophic complexity, dispersal, and stochastic disturbances—alongside the Nature Contributions to People (Regulatory, Material, and Non-material) to develop three distinct scenario narratives (Table 1, S2). This process was conducted in collaboration with the Rewilding Oder Delta (ROD) team and an expert panel of researchers.

**TABLE 1.** Key elements from the overall scenario storylines. The table outlines the key elements of rewilding and the Nature Contributions to People as described in the narrative scenarios. (see Table S2 for detailed information).

|  |  | Scenario |  |  |
| --- | --- | --- | --- | --- |
|  |  | Nature for Nature<br>(NfN) | Nature for Society (NfS) | Nature as Culture - One<br>with Nature (NaC) |
| Main narrative |  | Nature comeback -<br>All rewilding actions<br>are centred on<br>restoring lost<br>species interactions<br>and ecosystem<br>functions. | Maximising nature<br>contributions to people-<br>rewilding to enhance<br>nature's benefits to people<br>through sustainable<br>management. | Living in harmony with<br>nature- Community-based<br>rewilding for local identity<br>and stewardship |
| Action | Component | Nature for Nature<br>(NfN) | Nature for Society (NfS) | Nature as Culture - One<br>with Nature (NaC) |
| Rewilding framework (Perino et al.,) | Trophic Complexity | Strict actions for habitat protection and keystone species rehabilitation (e.g., nesting programs) | Reintroduction of species with a focus on restoring ecosystem services (e.g., pest regulation through predation and pollination). | Restoring trophic complexity through return of emblematic species (e.g., Bison, Elk and Grey seal) |
|  | Connectivity | Enhancing connectivity with green and blue corridors to link protected and high biodiversity value areas | Promoting connectivity through adaptive management practices. | Promoting connectivity for diverse and heterogenous biodiversity-friendly landscapes |
|  | Natural Disturbances | Rewilding efforts focus on rewetting dried peatlands to natural stochastic dynamics and biodiversity conservation. | Restoring natural disturbances to improve water and soil quality in peatlands | Community-based rewetting management for natural disturbances and biodiversity-friendly practices |
| Nature contributions to people | Regulatory services | Regulation of freshwater quality through the restoration of riverbeds and flood plains | Regulating freshwater quantity and flooding regimes through adaptive management | Climate regulation through the restoration of peat soils for CO2 absorption |
|  | **Material Services** | Reduce intensity of farming and wood extraction.<br>Improvement of water resources. | Intensive farming through precision agriculture;<br>sustainable wood extraction | Organic farming and paludiculture g for renewable energy and construction materials |
|  | **Non-material services** | Improvement of well-being through the creation of new spaces for enjoyment of natural landscapes | Improving the local economy through sustainable management and technology in agriculture | Landscape stewardship, community-based rewetting, and youth engagement for biodiversity-friendly farming |

### 2.5 Scenario mapping

To capture stakeholder perspectives, we conducted one-hour semi-structured online interviews via videocall, structured in four phases: rapport-building, introduction to the scenario storylines, participatory mapping, and discussion of preferences, co-benefits, and trade-offs (Table S5). Public Participatory GIS (PPGIS) is utilised to gather stakeholder preferences for future land use by enabling participants to visualise landscape changes through three storylines (Fagerholm et al., 2021). During participatory mapping, Stakeholders were asked to draw on the map to illustrate how specific land use types would change over a 30-year period. The participants were encouraged to identify areas of interest, highlight potential conflicts, and suggest preferred rewilding actions on the provided maps. Each land use type was represented in different colours to facilitate the reporting and analysis of the results.

Afterward, participants held in-depth discussions to express their preferences and examine the associated trade-offs and co-benefits of each action. For specific questions and the maps used during the interviews, refer to Table S5 for a detailed overview of the interview structure. Finally, Stakeholder inputs are recorded using GIS tools and digitised maps, and qualitative insights were documented through facilitated discussions. The outputs from the PPGIS process were used to develop heat maps that indicates priority zones for rewilding based on stakeholder input.

### 2.7 Qualitative Data analysis

We conducted a systematic content analysis, combining qualitative and quantitative methods, to categorise and cluster transcribed interview data (Mayring, 2014). Content analysis, a widely used tool in social sciences, allowed us to systematically collect, summarise, and categorise data through coding, categorisation, synthesis, and frequency analysis. The Interview data was organised into a matrix to identify key themes related to rewilding and Nature’s Contributions to People (NCPs), categorising them as actions. We used a mixed content structuring approach, integrating both deductive and inductive category formation to ensure a thorough synthesis of insights.

In the PPGIS processes, stakeholders marked their preferences on a current land-use map, using colour codes for different land-use types. These maps were then digitised into GIS software, clustering similar land-use types denoted by stakeholders into four categories: protected area, reflooded area, agricultural land, and green-blue corridor. The PPGIS results were analysed in two steps: identifying spatial patterns of landscape values and clustering them into a heat map highlighting areas deemed most valuable for rewilding. This approach enabled a comprehensive visual and quantitative analysis of potential land-use changes based on stakeholders’ values and preferences.

## 3 RESULTS

The outputs from the stakeholder interviews were instrumental in informing participatory mapping and assessing perceptions of land-use changes. The findings are organized into two main sections. First, a systematic content analysis of the interview data was conducted, creating a comprehensive list of rewilding actions linked to each scenario. This analysis highlights the diverse perspectives and priorities of the stakeholders, categorizing these actions by scenario. Second, the spatial aspect of these actions was examined through participatory mapping, analysing their distribution and patterns. This spatial assessment provides insights into stakeholder preferences for the placement of rewilding initiatives and identifies potential synergies or conflicts arising from differing land-use visions.

### 3.1 Identification of rewilding actions

All scenario narratives agreed on the need for actions restoring natural disturbance dynamics and bolstering landscape connectivity through measures like river and peatland reflooding (Fig 2).

Under a Nature for Nature scenario (NfN), the resulting narratives prioritised connectivity and natural disturbance restoration to increase wildness; the focus is on protected and other ecosystems under low human resource use. Overall, stakeholders identified the creation of blue and green corridors as a primary target of rewilding, with two benefits: facilitate free movement of species and restore the region’s long-lost water dynamics (Fig 2A). Furthermore, stakeholders coincided in prioritising actions aimed at achieving self-regulated ecosystem processes both in terrestrial and freshwater realms. These included halting wood extraction in some forests, the recovery of more natural groundwater dynamics, and reflooding dried peatlands. A smaller group of stakeholders mentioned actions to restore trophic complexity in the region, notably breeding and feeding habitat restoration programs for the lesser spotted eagle (Fig 2A). Some indirect benefits of rewilding to people were the provision of better water quality and quantity and carbon sequestration in peatlands (Fig 2A-NCPs section).

Under a Nature for Society scenario (NfS), stakeholders chose actions to reduce land use intensity by implementing technological advances. Similar to the previous scenario, rewilding was associated with restoring natural processes, e.g., allowing for natural succession in grasslands and rewetting drained peatlands not used for agriculture. However, the main emphasis was on the sustainability of material contributions to people through regulating wood extraction and biodiversity-friendly agriculture, such as adopting precision agricultural production to reduce pollutants (Fig 2B). Similarly, non-material nature contributions, such as participatory flood delimitation, were actions often mentioned by the interviewed stakeholders. Other important but less frequently mentioned actions include regulatory services such as flood regulation and water dynamics regulation. In this scenario, protected areas primarily aim to restore grasslands and forests by actively harnessing natural succession in non-productive lands.

In the “Nature as Culture” (NaC) scenario, stakeholders emphasised the need to implement co-production and community-based actions. Rewilding initiatives aimed to promote landscape stewardship, support local economies through biodiversity-friendly agriculture and reintroduce emblematic species to enhance local identities and cultural value. These initiatives often mentioned regenerative agricultural practices, such as sustainably harvesting reeds through paludiculture to provide raw construction materials (see Fig 2C). The potential benefits of paludiculture for water quantity provision and carbon sequestration were favoured.

**FIGURE 2.**
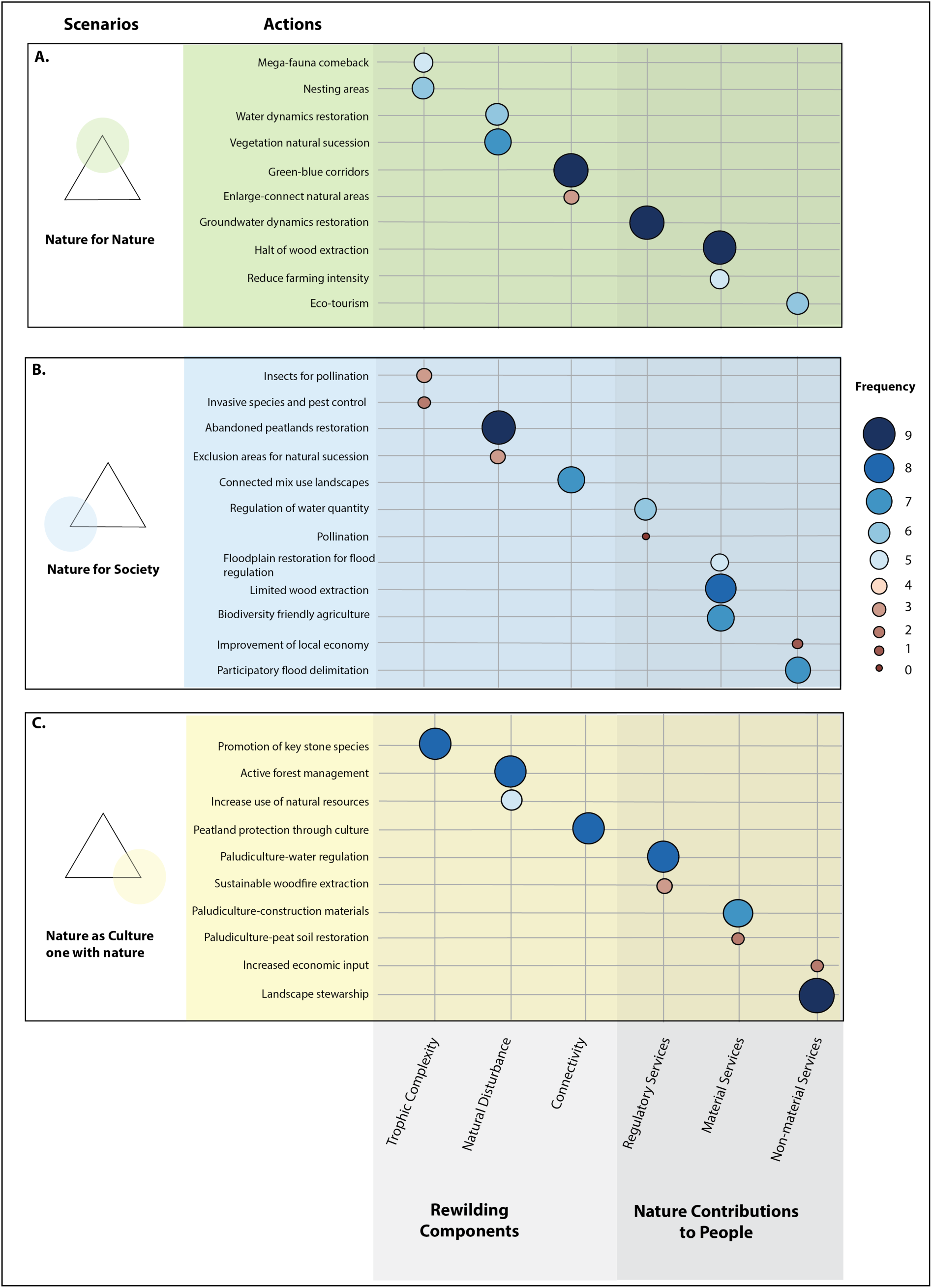
Frequency of main actions mentioned by the stakeholders for the different NFF scenarios. The circles represent how often specific actions were mentioned in different scenarios. The size and colour of the circles indicate how often these actions were mentioned. Smaller red circles represent actions mentioned less often, while larger, darker blue circles represent actions mentioned more often.

### 3.2 Mapping Scenario Storylines

The stakeholders’ mapping of all scenario actions resulted in key priority areas for rewilding and land use management under the different scenarios. After assembling the spatial priorities, we grouped actions into four major categories: expansion of protected areas and management changes, reflooding, creation of green and blue corridors, and biodiversity-friendly agriculture (Fig 3).

**FIGURE 3.**
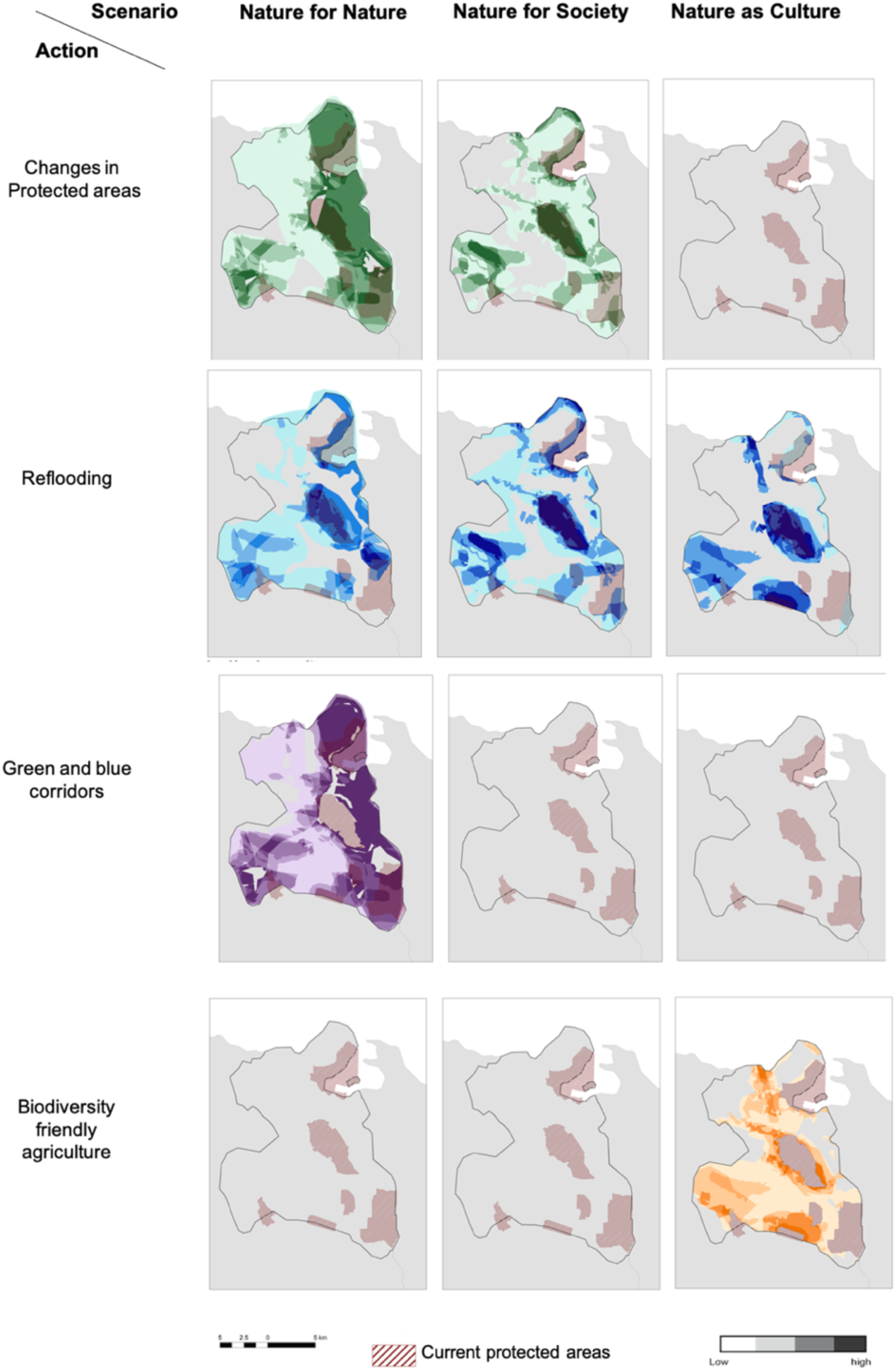
Heatmap of rewilding actions by scenario, as identified after overlapping the areas of interest identified by the stakeholders. Darker colours represent areas identified by a larger number of stakeholders.

The NfN scenario mapping resulted in large rewilding areas covering most of the study area (Fig 3A). Most of the actions took place along the border with Poland, an area characterised by extensive coniferous and broadleaf forests. Key rewilding outcomes include higher forest and transboundary ecological connectivity, a larger old-growth forest cover, and the recovery of functional peatlands. Connectivity in wetlands is to be restored through dike and dam removal, resulting in large, connected freshwater habitats. The expansion of protected areas was prioritised in roadless areas. With these actions, an increase in connectivity across freshwater and terrestrial realms would benefit the passive restoration of the trophic complexity.

In the NfS and NaC scenarios, most actions aimed to de-intensify land management, although there were no major land-use changes. The NfS scenario extended protected areas, but these were smaller than in NfN and were allocated as buffer zones at the interface with farming and forestry areas (Fig 3B). Reflooding is prioritised in peatlands and grasslands along rivers and coastal areas to protect farmland and urban areas against floods. As a result, highly heterogeneous landscapes would dominate, shaped by diversified uses.

The NaC scenario transformed intensive agriculture areas into extensive forms, with widespread implementation of reflooding. Protected areas did not increase. Instead, reflooding was allocated in some farming areas to create new semi-natural habitats compatible with extraction activities (e.g., paludiculture). Areas surrounding croplands, grasslands, and rivers were designated for rewilding, creating a landscape mosaic.

When considering only the areas of interest mapped by five or more stakeholders (marked as darker green in Fig 4.A), the total area allocated to NfN actions covered 65.8%, followed by NfS (= 14.4%) and NaC (= 8.6%). By converging the layers of the three scenarios on top of each other, we end up with a map that shows areas where two or three scenarios converge or areas where only one type of scenario is found (Fig 4B). For areas where only one type of scenario is found the narrative of NfN exclusively covered 47.2% of the landscape, followed by NaC with 2.36% and NfS with a total of 0.16% of area covered.

**FIGURE 4.**
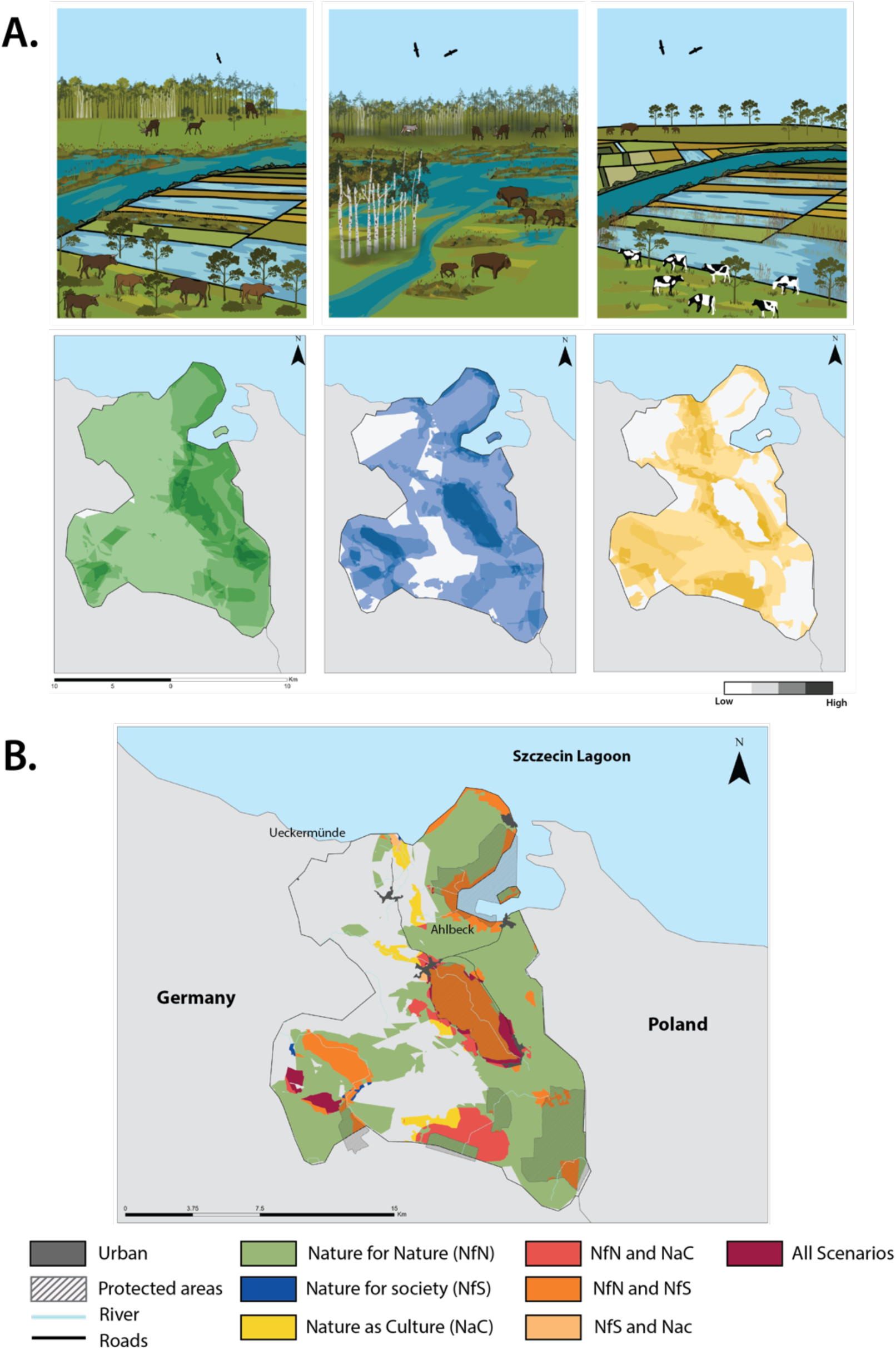
A. Overall change in land use considering the different scenarios B. Overlap of the areas identified by stakeholders as holding potential for more sustainable practices under the different nature future scenarios.

Regions where the NfN and NfS scenarios overlapped encompassed 14.06% of the study area (Fig 4B). Actions in these regions included strategies such as reflooding for disaster risk management and innovations to support biodiversity-friendly practices. The overlap between NfN and NaC, which only covered 5.6% of the total landscape, involved zones for transitioning to small-scale agricultural practices, which resulted in material provisions such as construction materials and food. The overlap between NfS and NaC covered 2.13% of the area (Fig 4B). These areas were distributed along traditional grasslands, agricultural fields, and towns. The main associated rewilding actions included the restoration of natural water dynamics and the improvement of agricultural yield.

Areas of interest for the three scenarios encompassed only 1.91% of the study area. These were distributed adjacent to natural parks, extensive livestock grasslands, and urban settlements. As demarcated by the stakeholders, these areas hold a high nature value in terms of biodiversity conservation and providing services such as water quantity and cattle feed production. These smaller areas are distributed along the landscape, creating a mosaic between all three narratives.

## Discussion

The Nature Futures Framework (NFF) has been effectively implemented on a large scale, as shown in studies by D’Alessio et al., (2024) and Diogo et al. (2025), focusing on nature-centred narratives in Europe. However, a gap exists in applying these narratives locally to actionable rewilding efforts. Our research addresses this by exploring participatory scenarios for diverse nature futures, revealing the benefits and trade-offs of rewilding practices. Our Findings indicate show that considering diverse natural values can enhance rewilding outcomes, highlighting a gradient of actions from passive to active rewilding for effective landscape stewardship.

### Identification of rewilding actions

By mapping rewilding components as foundational elements for developing three scenario narratives, we were able to develop a synergistic approach by addressing the shortcomings of past participatory scenarios that often focused solely on either instrumental or intrinsic values. In our process, we encouraged stakeholders to identify actions that balance these values without prioritising one over the other. This approach allowed us to move beyond seeing rewilding only as a nature-for-nature effort (Dunn-Capper et al., 2023; Quintero-Uribe et al., 2022). Our findings suggest that, unlike previous scenario exercises that unintentionally guided stakeholders toward land management actions favouring instrumental values, rewilding components such as improving landscape connectivity and reflooding peatlands were transversal across all three scenario narratives. However, the specific management actions varied depending on the underlying value framework, reflecting the diverse needs and priorities of the local community.

In the “Nature for Society” scenario, stakeholders supported a mix of passive and active rewilding strategies, such as peatland restoration, river reflooding, and forestry reduction, to enhance landscape connectivity and natural disturbances (Torres et al., 2018). These actions were primarily driven by their benefits to ecosystem services like water regulation and carbon sequestration (Couwenberg et al., 2011). Similarly, the “Nature for Nature” scenario prioritised improving connectivity and restoring disturbance regimes, reconnecting fragmented landscapes, facilitating the movement of species, and enhancing biodiversity with a focus on self-sustaining ecosystems and biodiversity conservation (Montes-Rojas et al., 2024). Despite this intrinsic value framing, these measures indirectly benefited people by enhancing water availability and quality. Also, in the “Nature as Culture” scenario, stakeholders favoured actions like peatland rewetting to support extensive agriculture (i.e. paludiculture) while restoring natural hydrological processes and reducing human intervention (Kreyling & Tanneberger, 2023).

The convergence of rewilding efforts across various scenarios can reveal important regional priorities for natural resource management and biodiversity conservation. Throughout our discussions with stakeholders, many highlighted the need to ensure long-term water resources to support agricultural productivity while safeguarding crucial ecosystems (Bauwe et al., 2012; Marx et al., 2023). They emphasised the need to tailor diverse rewilding actions to address future challenges and foster resilience within the community. This situation reflects ongoing social and environmental pressures and the importance of adaptive land management strategies that can effectively respond to future changes.

Stakeholders predominantly associate the reintroduction of keystone species with cultural values. This perspective differs from previous studies, which often emphasised the intrinsic (NfN) or instrumental (NfS) value of trophic rewilding (Schmitz et al., 2023; Svenning et al., 2016). Thus, highlighting the need for rewilding strategies that incorporate cultural aspects alongside ecological and economic considerations. Our findings reinforce existing literature that underscores the importance of involving local communities in rewilding efforts. Increased engagement with local communities has been identified as a crucial factor in fostering public acceptance of rewilding initiatives, particularly in relation to human-wildlife interactions and potential conflicts (Segar et al., 2022).

Understanding the connections between intrinsic, instrumental, and relational values aids in designing land management strategies that meet conservation goals and address local community needs (Fischer et al., 2020). Future research should look into how participatory scenario approaches can help shape policies and create more inclusive and flexible conservation strategies.

### Mapping of Scenario Storylines: Spatial allocation of rewilding actions

In addition to identifying potential rewilding actions, we employed participatory methods to assess co-benefits and emerging land-use challenges. Using a PPGIS approach to represent rewilding actions spatially, we analysed their distribution, revealing key trade-offs and synergies (Kyem, 2021). Heat map analysis of the three scenarios highlighted areas with strong stakeholder consensus for implementing specific rewilding strategies, providing valuable insights for future land management.

The mapping results indicate that 48% of the identified rewilding areas align with a “Nature for Nature” (NfN) perspective. It corresponds with regional management plans that aim to transition large patches of forested land from commercial forestry to protected areas, abandoned sites, or reduced timber harvesting to facilitate the development of old-growth forests over the next 40 years. Some privately owned areas already serve as pilot sites, undergoing gradual reductions in forest management to support this transition (German Federal Environmental Foundation, 2023).

Smaller and more dispersed areas were allocated exclusively for “Nature as Culture” (NaC) scenarios, creating opportunities for biodiversity-friendly agriculture and biocultural conservation. Actions such as peatland rewetting were proposed to support smallholder farming while restoring natural hydrological processes (Durán et al., 2023; Van Meerbeek et al., 2019). For example, stakeholders identified priority areas for paludiculture—agricultural practices on rewetted peat soils that minimise environmental impact while enabling peat restoration. Biomass harvested from native wetland vegetation, such as reeds and sedges, can provide raw materials for biofuels, insulation, and construction (Tanneberger et al., 2020). This approach supports local livelihoods and enhances long-term ecosystem resilience by maintaining stable water tables and natural disturbance dynamics (Harvey & Henshaw, 2023; Torres et al., 2018). Beyond direct ecological benefits, rewilding in NaC areas can also help restore human-wildlife relationships (Schmölcke & Grimm, 2024). For instance, reducing pressures on forest ecosystems along the German-Polish border could facilitate the return of charismatic species such as bison, elk, and lynx (Kowalczyk et al., 2013).

### Managing Land-Use Conflicts and Synergies

While certain areas align with a single value perspective, others exhibit overlapping interests where rewilding can support multiple objectives. Transition zones between protected areas, agricultural land, and urban spaces are significant, encompassing diverse ecological, economic, and cultural values (Coles et al., 2018; Mann et al., 2018). However, these areas also present potential land-use conflicts, as multiple stakeholders may have competing interests. Finding a balance between conservation and land use requires measures that integrate ecological restoration with local socio-economic needs.

For example, peatland restoration efforts must consider hydrological recovery and stakeholder engagement. Maintaining natural water levels can enhance ecosystem function, but successful implementation requires collaboration with local communities. Engaging landowners, providing financial incentives, and integrating rewilding into local land-use planning is crucial to long-term success (Ziegler, 2020). Similarly, species reintroduction programs must incorporate compensation schemes and promote non-lethal predator control techniques to gain public support (Killion et al., 2020; Pedroza-Arceo et al., 2022).

Certain areas also present opportunities to integrate conservation with sustainable land management. Overlaps between NaC and NfS scenarios highlight the potential for biodiversity-friendly agricultural practices that enhance food production while improving water quality and climate resilience (Tanneberger et al., 2022; Ziegler, 2020). Rewilding actions that promote the co-production of ecosystem services, such as flood mitigation and drought resilience, can help mitigate climate change impacts while maintaining ecosystem integrity (Harvey & Henshaw, 2023; Schmitz et al., 2023; Svenning, 2020; Temmink et al., 2023).

One promising approach is the adoption of paludiculture, which cultivates wet-tolerant plants on rewetted peatlands. This method generates economic benefits through biomass production while simultaneously improving water purification, nutrient retention, and carbon sequestration (Joosten et al., 2016; Kreyling & Tanneberger, 2023). Furthermore, paludiculture supports peatland-dependent species, contributing to biodiversity conservation (Bockermann et al., 2023; Tanneberger et al., 2022). The mapping methodology used in this study provides a valuable tool for identifying and anticipating land-use conflicts. By integrating stakeholder perspectives and scenario-based approaches, our framework highlights areas where trade-offs between ecological restoration and socio-economic activities may arise. This enables policymakers to proactively address conflicts and align rewilding initiatives—such as paludiculture—with broader sustainability goals.

### Benefits of Participatory Scenario Planning for Rewilding

Participatory scenario planning (PSP) offers a structured way to integrate diverse stakeholder perspectives into rewilding efforts, balancing ecological restoration with social and economic considerations. By fostering dialogue and identifying trade-offs, PSP enhances adaptive decision-making and local acceptance of landscape transformations (Killion et al., 2020; Quintero-Uribe et al., 2022), particularly in Europe, where competing land-use demands and conservation priorities require careful coordination (Perino et al., 2019). Implementing community-based management alongside rewilding can mitigate conflicts, such as flood risks in cattle-grazed peatlands. Transitioning farmers to alternative practices, like peatland biomass cultivation, through training and financial support helps align ecological goals with local livelihoods (Wichmann & Nordt, 2024).

Proactive community involvement also benefits species reintroductions, such as bison and elk. Educating local populations on the ecological benefits of species restoration and promoting non-lethal coexistence strategies can reduce human-wildlife conflicts and foster long-term support (Carter & Linnell, 2023; König et al., 2020). PSP aids this process by enabling stakeholders to explore different future scenarios, anticipate challenges, and co-develop solutions.

Participatory mapping results can improve land-use planning by pinpointing high-impact rewilding actions across scenarios, such as peatland reflooding, which enhances connectivity and aligns with regional conservation strategies. This ensures efficient resource allocation in overlapping scenario areas (Metzger et al., 2017; Peterson et al., 2003). We strategically selected nine stakeholders knowledgeable to Nature Conservation decision-making. While a small group can limit diverse viewpoints and may introduce bias (Pahl-Wostl, 2002), it can also lead to valuable insights in the early stages of a project. By carefully selecting participants and using structured methods like PPGIS, we ensured the reliability of the information we collected. Focusing on depth instead of breadth helped us explore future possibilities in rewilding. This helped us understand the region’s complex ecological and social dynamics (Quimby & Beresford, 2023; Rohrbach et al., 2016). Our findings show that participatory mapping and respondent-driven sampling can produce reliable and useful data.

Future efforts will focus on re-engaging local communities to share the outcomes of participatory mapping and discuss rewilding actions’ practical implications. Broader stakeholder engagement will refine scenario narratives and improve calibration of rewilding strategies while addressing socio-economic constraints. Integrating the Nature Futures Framework (NFF) into rewilding can help illuminate complex socio-ecological dynamics and maximize co-benefits, despite some limitations in recognizing socio-economic constraints (Dunn-Capper et al., 2023). By employing PSP in rewilding frameworks, policymakers can better navigate socio-ecological complexities and enhance the long-term benefits of conservation initiatives.

### Developing a Framework for Rewilding landscapes

This study introduces a methodological framework for creating participatory scenarios to inform rewilding initiatives. By incorporating participatory processes, we offer a structured approach to pinpointing priority areas for rewilding, while also addressing the socio-ecological complexities associated with landscape restoration. Our framework presents a replicable methodology suitable for various contexts, serving as a practical tool for investigating rewilding and restoration efforts through the perspective of the Nature Futures Framework (NFF).

Moreover, we illustrate the practical value of structured methodologies in land-use planning, particularly through participatory mapping (Balram et al., 2004; Lim et al., 2021). These techniques enable decision-makers and practitioners to prioritize rewilding initiatives with the greatest potential impact across various scenarios. By spatially allocating scenario narratives alongside their related rewilding actions, our methodology serves as a valuable decision-support tool. It allows land managers to visualize potential land-use changes and pinpoint critical areas where multiple values and interests intersect. This approach enhances the ability of practitioners and policymakers to effectively target rewilding interventions, ensuring alignment with the specific ecological and societal needs of a given region.

Beyond its immediate application, our approach enhances broader decision-making processes by incorporating a range of values linked to nature (Fischer et al., 2020; Perino et al., 2022). This study demonstrates how participatory methods can be integrated into scientific and governance frameworks to foster more inclusive and adaptable rewilding initiatives. By connecting rewilding efforts with societal well-being and sustainable development goals, we highlight the significance of these approaches. The insights derived from our research provide a model for rewilding initiatives beyond the Oder Delta region, contributing to a more comprehensive and actionable framework for landscape restoration.

## Supporting information

Supplementary material linked to the methodology of participatory scenario planning

## Acknowledgements

The authors gratefully acknowledge the support of iDiv funded by the German Research Foundation (DFG-FZT 118, 202548816). LQ-U was supported by funding from the European Union’s Horizon 2020 Programme “Marie Sklodowska-Curie Innovative Training Networks” (MSCA-ITN-ETN, grant no. 813904), and from the FEdA programme of the Federal Ministry of Education and Research of Germany (BMBF) (grant no. 16LW0064K).

We thank Jennifer Hauck from CoKnow Consulting for her early input during the conceptualization and design of the work, and the Rewilding Oder Delta e.V. for their advice and support in different study stages.

## Notes

### Competing Interest Statement

The authors have declared no competing interest.

