## Supplementary material linked to the methodology of participatory scenario planning for "Finding space for rewilding: participatory scenarios reveal ecological opportunities based on plural values of nature"

FIGURE S1. FLOWCHART OF SCENARIO DEVELOPMENT STEPS

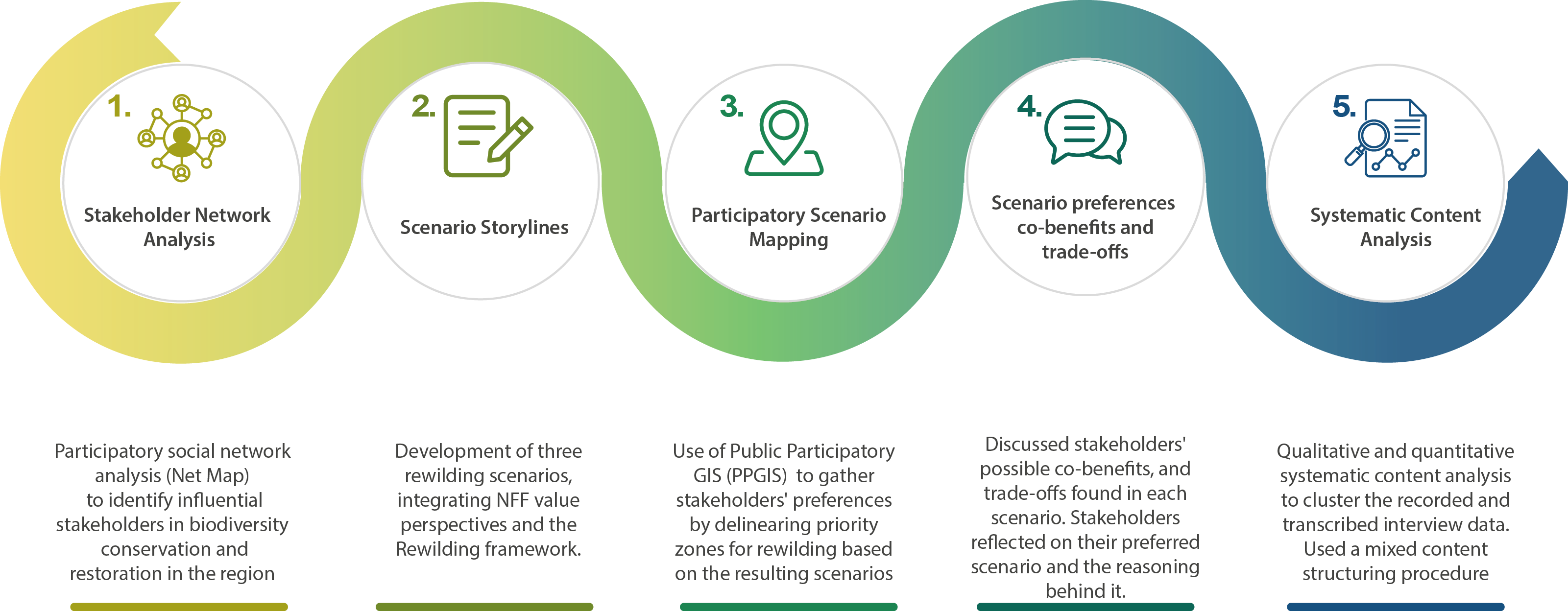

TABLE S2. DETAILED SCENARIO STORYLINES

| **Nature for Nature- Rewilding for Nature Comeback** | **Nature for Society - Rewilding for maximising Nature Contributions to People** | **Nature as Culture*:* Rewilding living in harmony with nature** |
| --- | --- | --- |
| The rewilding actions actively focus on restoring lost species interactions and ecosystem functions, intending to reduce landscape management and preserve nature's diversity and functions, enabling the restoration of self-sufficient and intricate ecosystems. It is accomplished by actively restoring these interactions and functions by expanding protected areas with stricter protection measures and promoting programs that facilitate the rehabilitation of keystone species, such as nesting programs, thereby enhancing trophic complexity. Connectivity is improved by creating green and blue corridors that connect protected and high biodiversity value areas, while also allowing a natural succession of vegetation in old-growth forests and abandoned areas. Finally, for restoring stochastic disturbances, the rewilding actions focus on rewetting dried peatlands to foster biodiversity conservation and restoration. | Enhance the benefits of nature by implementing sustainable management practices that promote biodiversity, a natural capital-based approach, and active management of natural resources. In this scenario, the rewilding actions centred on providing and restoring different Nature Contributions to People by increasing trophic complexity through the reintroduction of key species, such as wolves and insects, that can provide essential ecosystem services such as pest regulation through predation and pollination. Furthermore, connectivity was promoted through the sustainable use of natural resources in forests, and grasslands by decreasing management intensity, thus improving connectivity and creating buffer zones between protected areas and other land uses. Finally, the restoration of natural disturbances was centred around providing regulatory services to reduce the risk of flooding and improving the quantity and quality of water and soil through the restoration of water dynamics in abandoned peatlands. | Prioritising and community-based management of natural resources, lifestyle changes, and education. This scenario promotes local identities and landscape stewardship by designing rewilding actions that reflect these values. The restoration of trophic complexity aimed to bring back emblematic species (e.g., Bison, Elk and Grey seal) that can foster local identities. Connectivity of landscapes is promoted to create a rich but heterogeneous landscape that fosters diverse biodiversity-friendly practices, enabling species to overcome natural barriers. Finally, the promotion of natural disturbances was achieved community-based rewetting management. An adaptive management approach was followed through seasonal reflooding of peat soils and grasslands. In areas with high potential for peatland restoration, biodiversity-friendly practices were promoted. Overall, this scenario emphasises the need for community-based management and highlights the cultural and social dimensions of rewilding. |

FIGURE S3. NETWORK MAP FOR BIODIVERSITY CONSERVATION STAKEHOLDERS IN THE ODER DELTA.

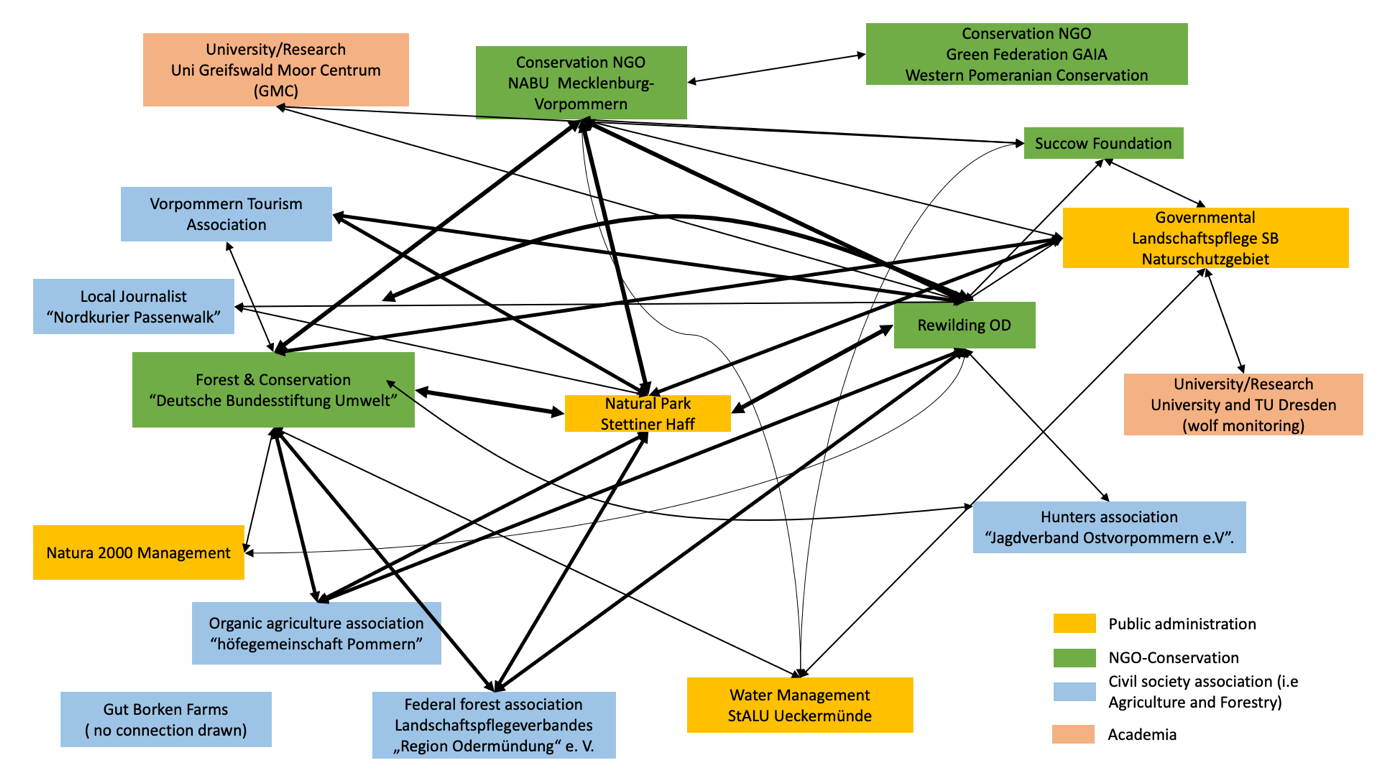

TABLE S4 OVERVIEW OF STAKEHOLDERS SELECTED FROM THE NET-MAP ANALYSIS

| **Netmap Releance**  **(Links)** | **Stakeholder Group** | **Working area** | **Contacted** | **Interviewed** |
| --- | --- | --- | --- | --- |
| 9 | Civil society association | eco tourism | YES | YES |
| 8 | Conservation NGO | forest conservation | YES | YES |
| 8 | Public Administration | Tourism -land management | YES | YES |
| 7 | Public Administration | Nature Conservation | YES | YES |
| 7 | Public Administration | Nature Conservation | YES | YES |
| 6 | Conservation NGO - Academia | water management | YES | YES |
| 6 | Conservation NGO | conservation | YES | YES |
| 4 | Public Administration | water management | YES | NO |
| 4 | Conservation NGO | water management | YES | NO |
| 3 | Civil society association | sustainable agricultre | YES | YES |
| 3 | Civil Society association | Forestry - hunter | YES | YES |
| 2 | Civil society association | hunting sector | YES | NO |
| 2 | Civil society association | hunter | YES | NO |
| 1 | Academia | research | NO | NO |
| 1 | Conservation NGO | Nature Conservation | YES | NO |
| 0 | Civil society association | farmer | NO | NO |

TABLE S5. PARTICIPATORY SCENARIO INTERVIEW PROTOCOL.

|  | **Step by step guide** |
| --- | --- |
| **Main objectives for the interview approach** | Gather spatially explicit information of the changes in the landscape associated with different scenarios- participatory mapping  Identification of co-benefits and trade-offs of different NFF scenarios |
| **Interview approach** | Interviewers are the principal researchers of the project  The aim is to conduct about 10 interviews, each lasting about 1 hour.  During this time one interviewer will ask the questions and lead the interviews, while the other takes notes and visualised results.  The interviews will be:   - semi-structured, qualitative interviews - held online or in person on the ground - recorded, and additionally detailed field notes will be taken - done with one actor each |
| **Leading questions:** | 1) Where and how would the three scenarios change the landscape? 2) Which preferences do interview partners have concerning scenario(s) or individual restoration/rewilding activities? 3) Which co-benefits can be related to the three nature futures scenarios? 4) Which trade-offs can be related to the three nature futures scenarios?   If there is time, the following questions can be covered:   5) What could be supporting policies or instruments to realize scenarios or rewilding activities while at the same time fostering co-benefits? |
| **Interview guideline** | **Phase 1: Introduction & building rapport**   - Brief introduction of how the interview will proceed - Have interview partners agree to the consent form and start recording - Short round of introductions: - Principal researcher introduction with the goal and approach of her PhD thesis: developing scenarios for nature restoration and conservation, and models to assess possible impacts of the different scenarios on changes in land use, biodiversity, and ecosystem services. - Introduction of translator and consultant. - Interviewee briefly introduces themselves ( Description of their background and projects associated to the Oder Delta) - Brief introduction to the objective of the interview:   We are presenting you with three different conservation/restoration scenarios at once. We would like to ask you to try to give spatially explicit information about the changes in the landscape associated with the different scenarios. For this purpose, we will show you a map of Uckermünde Heide on which you can draw the changes. It is important that when assigning changes in the landscape the stakeholder imagine what could happen in 30 years into the future and push away from current political conditions.   - Introduce the three scenarios and allow questions of clarification (pay attention to not start to discuss the scenarios but only give clarifications!) |
|  | **Phase 2: Introduction of the Scenarios:**  Presentation of the three scenarios  **Scenario 1:**  This scenario is centered towards the creation or expansion of already existing protected areas. The delimitation of protected areas follows a stricter regulation where the intensity of management of the area is reduced and extraction of natural resources is only allowed outside the protected areas. This is done to foster biodiversity conservation and the restoration of ecosystem functions. The main actions that take place in the scenario is the prioritization of areas for restoring old grow forest to foster the recovery of species (and nesting programs), the rewetting of grasslands and peatlands with high potential for biodiversity restoration and the creation of green corridors to connect fragmented patches of protected areas.  **Scenario 2:**  Natural reserves are expanded or newly created in this scenario. However more land uses are allowed in the protected areas such as the sustainable logging and different forms of organic farming should be able to take place around the protected areas. Species that play an important role in pollination and control of pests are prioritized when restoring the landscape. Additionally, rewetting take place along rivers and forest to restore the natural floodplains.  **Scenario 3:**  In this scenario local farmers are encouraged to transition towards more environmental and biodiversity friendly practices that foster rich and diverse agricultural ecosystems. Natural protected areas remain the same in this scenario. The reinforcement of local populations of the grey seal, the lesser spotted eagle and the elk helps the local population to create a strong link with the identity of the area thus becoming in the long-term emblematic species of the Oder Delta. Finally, the rewetting of grasslands and peatlands only takes place where there is a high potential for biodiversity restoration and it does not result in the loss of grasslands for productive purposes |
|  | **Phase 3: Participatory Mapping**   - The interviewees will be presented with a map that has information of current land uses of the area (i.e protected areas, peatlands moores, grasslands and agricultural sites). They will be asked to indicate either if they think the area of the already existing land use category can increase or a new area for the land use should be created in the next 10-30 years. It is important to highlight that we want information at the local level. - The mapping exercise will be done for each scenario separately. However we based on previous test we group the questions per scenario in the following categories:  1) Increase or expansion of protected areas? 2) Where organic agriculture or forms of sustainable agriculture could take place 3) Rewetting of peatlands and moors 4) Creation of green corridors  - Before starting the interview is important to contextualize the stakeholders: First we want information at the local level and second, we will tell them to imagine changes ignoring current land ownerships. - Start of the participatory mapping process:  1) Protected areas/ecological agriculture.   In two of the three scenarios, protected areas would be expanded or newly created with two different categories of protected areas: in scenario 1, land use (agriculture and forestry) would be stopped, but people could still go there (e.g. for tourism use). In scenario 2, organic agriculture would be allowed to take place in protected areas. Where would agriculture take place in these areas?  Where would they draw in the protected areas in the first two scenarios (different areas or the same areas)?  In the third scenario, no protected areas would be expanded, but people would voluntarily practice organic farming. Imagine half of the agricultural land would be organic, where would this land be most usefully placed for biodiversity?  In scenario 1:  Q1a: Where would Strict protected areas be expanded or newly designated?  Q1b: Where would old, unused forests increase?  In scenario 2:  Q2: Where would organic agriculture/ sustainable logging could increase in the protected areas? Or outside?  In scenario 3:  Q3: 50% of Cultivated areas transition towards organic agriculture, where would this land be most usefully placed for biodiversity?   2) Rewetting:   Across all three scenarios, rewetting should occur but for different reasons.  In scenario 1  Q1: Rewetting of areas is to promote biodiversity and reinforce local populations. Where this would happen in the map?  In scenario 2  Q2: Rewetting of forest, peatlands and river beds to create natural floodplains, and improve the regulation of water dynamics of the area. Where this would happen in the map?  In scenario 3  Q3: The goal is to lose as little of productive grassland as possible during rewetting. Where this would happen in the map?  Do you see differences in the potential for rewetting on the map for the different reasons?   3) Green corridors   Q1: In Scenario 1, where on the map do you see opportunities to create ecological corridors? (explanation of green corridors: to connect separated patches of nature reserves) |
|  | **Phase 4: Scenario preferences**   - After mapping the stakeholder will be asked to state their preferences towards the three scenarios - If there is not an explicit preference, they are allowed to create a 4^th^ scenario with the components they like the most from each scenario.   Q1: Would you have a preference for a particular scenario, or which scenario would be the least desirable? |
|  | P**hase 5: Co-benefits and trade offs**   - After mapping the stakeholder will be asked to name the main co benefits for each scenario and if the different scenarios share co-benefits or trade-offs - It is important to give stakeholders a short explanation of what co benefits and trade-offs mean. (note: we should give the same example to all interviewees)   Q1a: Which co-benefits can be related to the different scenarios?  Q1b: Are there co-benefits shared between the scenarios?  Q2a: Which challenges can be related to the different scenarios?  Q2b: Are there challenges shared between the scenarios?  The answers will be recorded and noted by the interviewers |
|  | **Phase 6: Wrap up**  Ask interview partner, whether he/she has any additional questions or comments  Ask interview partners whom we should additionally interview  Ask if they are interested in keeping in contact with the project and if they are interested in seeing the outcomes of the interviews.  Thank interviews partners and end the interview. |
| **Documentation** | Documentation using online whiteboards; Audio recordings, detailed field notes, post script (the two interviewers meet immediately after the meeting and share their most important observations) |
| **Data analysis** | Qualitative content analysis, basic descriptive statistics |
